# Highly Dynamic Population of Owned Dogs and Implications for Zoonosis Control

**DOI:** 10.1101/2025.04.23.650047

**Authors:** Elvis W. Diaz, Stephanie Sila, Brinkley Bellotti Raynor, Micaela De la Puente-León, Michael Z. Levy, Ricardo Castillo-Neyra

## Abstract

**Background:** Domestic dogs are known carriers of a variety of zoonotic pathogens significant to public health including leishmaniasis, echinococcosis, and, the most lethal, rabies. Control efforts for rabies and other diseases focus on the removal of infected individuals and mass treatment or vaccination to reduce incidence and raise immunity in the susceptible population. However, the complex population dynamics surrounding free-roaming dogs in many developing countries complicate these control methods. While rabies vaccination coverage of 70% or higher, of the dog population at risk, performed annually, should be sufficient to maintain coverage above the critical threshold, rapid population turnover and growth could reduce effective coverage. Our objective was to investigate the impact of dog population dynamics on rabies control activities in a rabies-endemic city.

**Methodology:** Using data from annual door-to-door surveys in urban and periurban areas, we collected household information (e.g., number of residents), respondent characteristics (e.g., education level), and dog-related data (e.g., vaccination status). These variables were analyzed to evaluate changes in the dog population and vaccination coverage. A compartmental Susceptible - Exposed - Infectious - Vaccinated (SEIV) mathematical model was utilized to estimate the effects of population dynamics on herd immunity against rabies.

**Findings:** We found that the highly dynamic dog population in Arequipa and its rapid expansion hampers herd immunity, counteracting the efforts made in yearly vaccination campaigns and leading to underestimates of the human to dog ratios (∼4:1 vs 6:1). Moreover, there are significant differences in dog population trends within urban and periurban areas indicating that different strategies or coverage vaccination targets might be needed. This study also introduces an important variable of interest: *transient dogs*, a population with extremely high turnover (less than one year), which causes an important drop in vaccination coverage that has thus far gone undetected.

**Conclusions:** Stabilizing the free-roaming domestic dog population by lowering both mortality and birth rates may help improve the efficacy of current mass vaccination programs. Additionally, the population dynamic parameters, on which we base our models, must be regularly refined to best represent the current status of disease risk and population immunity. By using a One Health approach we can understand and address the dynamics of dog populations, and better control zoonotic diseases like rabies, ultimately improving public health.

## INTRODUCTION

Dogs harbor various pathogens, many of which can be transmitted to humans, causing zoonoses^1^. These zoonoses range from diseases with a relatively low impact on public health, such as scabies, to those with high mortality rates in both animals and humans, such as rabies^2^. The effectiveness of public health programs aimed at controlling dog-associated zoonoses can be significantly impacted by the dynamics of dog populations^3,4^, particularly free-roaming dogs, as this population is at highest risk of infection and most likely to transmit zoonotic agents to humans^3,5^.

Many developing countries, especially in Latin America, have large free-roaming dog populations^1,5,6^, which can carry various zoonotic pathogens^7^, *Rabies lyssavirus* being the most lethal. This virus, causing rabies, is almost always transmitted to humans through the bite of a rabid dog^8^. Consequently, elimination of human rabies, and other zoonoses, hinges on dog surveillance, the quick removal of infected individuals from the dog population, and mass dog rabies vaccination^9,10^, embodying a true One Health approach. Achieving dog vaccination coverage above 70% of the estimated at-risk dog population^11^ is crucial to interrupt transmission and reduce the circulation of the virus by reducing the population of susceptible hosts^12^.

Free-roaming dogs are more likely than restrained dogs to acquire and transmit the rabies virus; due to the higher contact rates^13,14^, and they are more difficult to handle and vaccinate^15,16^. If infected, these dogs can spread the rabies virus, or other pathogens, over longer distances^13,17^. Additionally, free-roaming behavior is associated with increased mortality and pregnancy rates^3,5^, which significantly influence dog population dynamics.

Accurate and regularly updated canine population studies are essential for planning rabies control programs ^3,9,18,19^. However, the dynamic nature of these populations poses challenges in obtaining precise parameters, necessitating continual re-evaluation to ensure effective rabies control.

While studies on estimates demographic parameters of dog populations in Latin America provide valuable insights, they often rely on cross-sectional studies^20^.Therefore, studies are needed to characterize highly dynamic populations taking into account *transient dogs*: populations that are very short-lived but large enough to impact public health interventions.

The city of Arequipa is experiencing an ongoing rabies epidemic with hundreds of dog cases detected since 2015. Here we present field data on the birth, mortality, and migration of owned dogs in Arequipa with the goal of updating our current model of rabies transmission and disease to better inform public health measures. Our objectives are to characterize the demographic structure of free-roaming dog populations, to compare the parameters of the canine population dynamics between urban and periurban areas in a large city of Latin America, Arequipa, Peru, and to explore the potential effect of the dog population turnover on mass dog vaccination campaigns (MDVCs) conducted to eliminate rabies in the area. Devising effective strategies to ultimately eliminate dog rabies and related human deaths can and should be informed by knowledge of the local dog population dynamics. This information may improve our understanding of how dog population dynamics differ across geographical areas, and how they influence the efficacy of rabies control efforts, such as vaccination coverage. Additionally, these findings can be applied to other dog-borne zoonoses to improve existing public health interventions (e.g., *Echinococcosis* treatment, xenointoxication for Chagas disease vector control)^21–23^.

## METHODS

### Ethics Statement

Ethical approval was obtained from Universidad Peruana Cayetano Heredia (approval number: 65369) and University of Pennsylvania (approval number: 823736).

### Study Setting

This study took place in the district of Alto Selva Alegre (ASA) within the city of Arequipa and one of the 29 districts of the province of Arequipa, Peru. The city of Arequipa is situated at around 2,300 meters above sea level and in 2017 had a human population of 1,092,744, of which approximately 87,570 live in the ASA district^24^. The rabies virus was first detected in the dog population in March 2015, with over 392 rabid dogs detected to date. At the moment this study was conducted, the canine population in ASA was estimated by the Ministry of Health (MOH) to be approximately 14,595 dogs, using a human-to-dog ratio of 6:1^25^.

The city of Arequipa spans a wide spectrum of urbanization, migration, and socioeconomic status. The city has a well define urban center, consisting of old established neighborhoods with a working network of sidewalks, roads, and access to transportation. In the outskirts of the city, there is a newer, periurban area. Compare to the urban center, the periurban area has a lower socioeconomic status, an increased crime rate and a more rugged, irregular terrain which makes travel more difficult. As a result, there are fewer community resources available to residents^26–28^. Both the urban and periurban areas are divided into localities, political subdivisions of the districts.

### Data collection

We conducted door-to-door surveys in both the urban and periurban localities immediately after the MDVC of 2016 and 2017. One adult, 18yo or older, was consented and surveyed in each household. The surveys were carried out with a web app created by our group for direct data entry on a mobile device. Household variables (number of people, highest education level, dog breeding), respondent variables (age, gender, education level) and dog variables (age, sex, vaccination status, number of puppies, degree of restriction, causes of dog absence in subsequent visits) were collected at each household.

In the first year of the study, 21 urban localities (4,531 houses) and 21 periurban localities (1,889 houses) were selected. In the second year, a subsample of the locations visited in the first year was selected. Four urban localities (1,066 houses) and 6 periurban localities (1,060 houses) were used for the second year’s sampling. All houses in the study localities were geocoded and the survey data were linked to the household coordinates. A new denomination termed ‘transient dogs’ was created to collect information on dogs that lived in the household in between surveys but was not alive at the moment of any survey.

### Statistical Analysis

Variables at the household and respondent level were analyzed such as dog ownership, gender, family size, and education level. At the dog level, variables such as age, sex, vaccination status, degree of restriction, and origin were analyzed. Comparisons were made of these variables between the first and second year of the study, as well as between the urban and periurban areas.

Categorical variables such as age, sex, and vaccination status were analyzed using a Chi-square test, or a Fisher’s exact test if statistically necessary (less than 10 observations). Discrete count variables such as the number of puppies per bitch were compared using a Poisson regression. The change in vaccination coverage was estimated by dividing the number of vaccinated dogs in 2016 that remained in the cohort by the MDVC 2017, by the total number of dogs present in the MDVC 2017. All statistical analyses, tables, and figures were generated with R (Team, 2013).

### Modeling the dynamics of the dog population and immunity against rabies virus

We estimated the population growth, the population turnover, the emigration and immigration rates, and the mortality and birth rates from the survey data. We assessed the impact of these population parameters on herd immunity using a compartmental mathematical model described in detail in Raynor et al, 2021^29^. Briefly, two different simulations of a basic Susceptible – Exposed - Infectious - Vaccinated deterministic compartment model were compared, one set with parameters from the periurban regions (higher population turnover) and one set with parameters from the urban regions.

## RESULTS

### Canine Demographics

We surveyed 673 houses in the urban area and 212 in the periurban area for a total of 885 houses (Table 1). Notably the mean number of dogs per house increased slightly between study years 1 and 2 in both areas (from 1.61 dogs/house to 1.72 dogs/house in the periurban area and from 1.44 dogs/house to 1.52 dogs/house in the urban area). Also, at the time of the first visit the density of dogs in the periurban geographic area was estimated to be 443 dogs/km2 and our data suggested that it increased by 8.7% over the course of one year. In the urban geographic area, the density of dogs was estimated to be 1,738 dogs/km2 and we estimated that it increased by 6.3% over the course of one year.

**Table 1.**
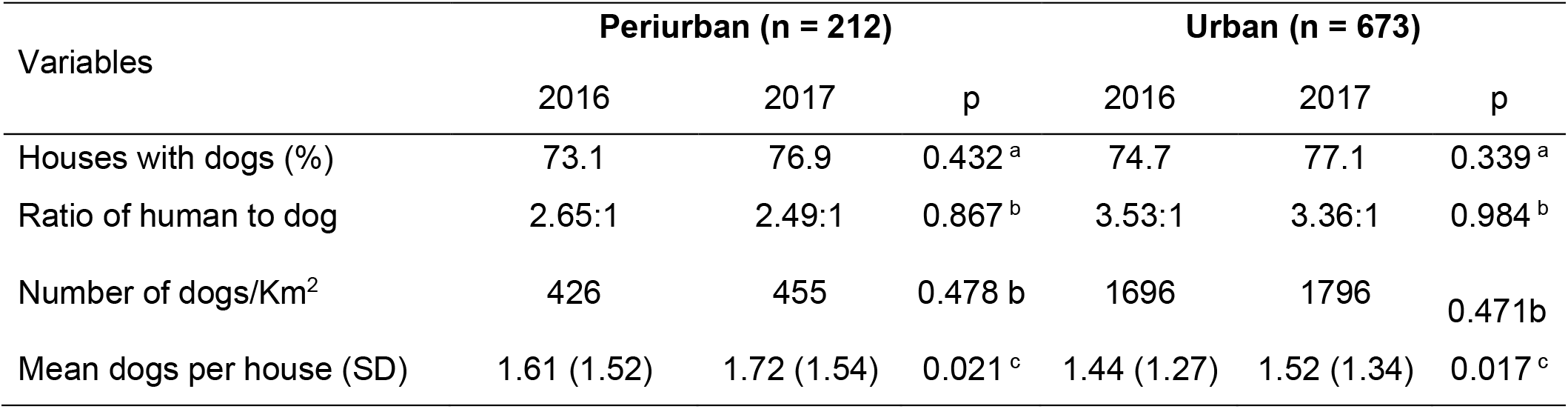
General information of surveyed houses. p values estimated with (a) Chi square test; (b) Bootstrap and (c) Student t-test

On average periurban dogs were younger than urban dogs (35.1 vs. 44.6 months old). The mean age of periurban dogs increased 2.3 months over a year (though not statistically significant), but the mean urban dogs’ age did not change. Females represented only a third of the dog population, with more females in the urban areas (36.9-37.8%) compared to periurban areas (27.9-31.1%) (Figure 2). No changes were observed in sex distribution between visitations. Dogs were more often reported as “purebred” in urban households (26.2-31%) as opposed to periurban households (7.9-14.1%).

**Figure 1.**
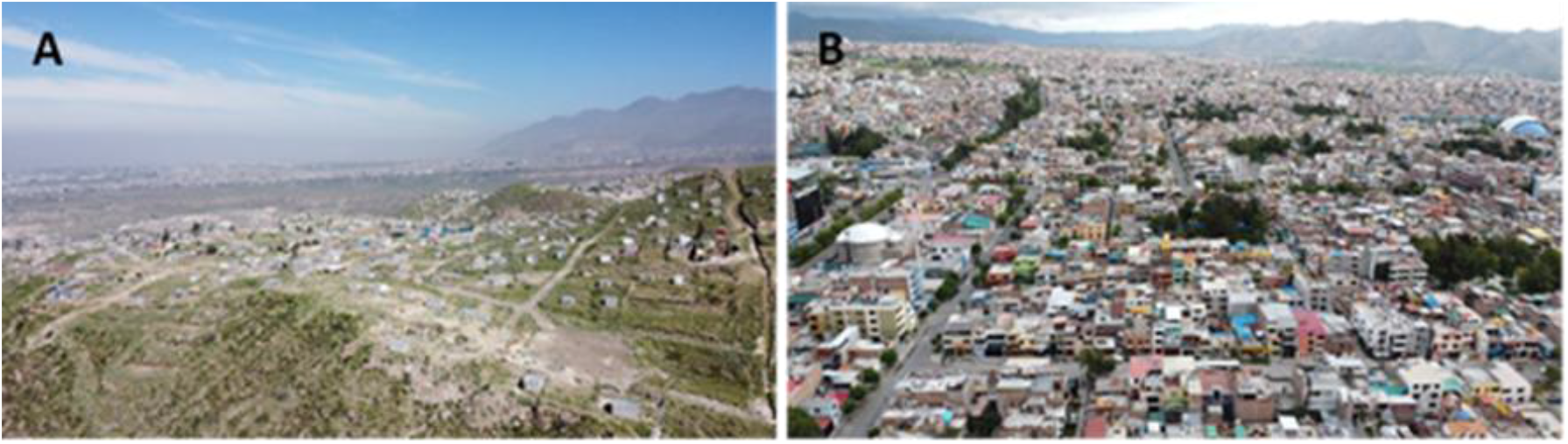
Locations of (A) Periurban, and (B) Urban study communities in Arequipa, Peru.

**Figure 2.**
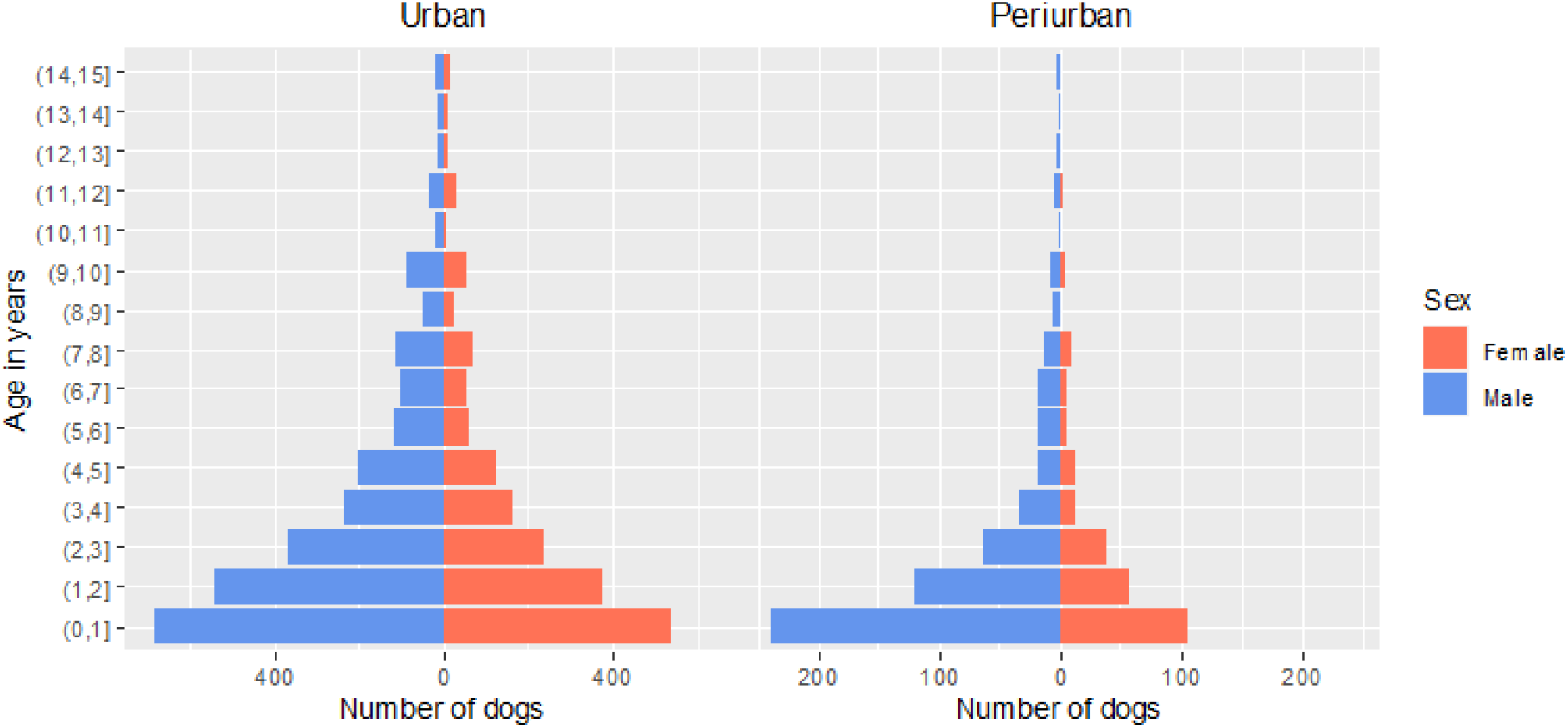
Baseline sex and age distribution in urban and periurban areas of Arequipa, Peru.

### *Transient dogs* are more prevalent in periurban areas and contribute to population turnover

Regarding *transient dogs*, dogs that remained at home for a short period between the first and second surveys, we observed a significant difference in the proportion of houses with *transient dogs* between periurban and urban areas. In periurban areas, 24.5% of the households surveyed reported having had *transient dogs*, compared to only 13.0% in urban areas (p=0.006; Table 4). We also found different causes for the presence of *transient dogs*. Death was the main cause for 66.7% of periurban transient dogs and for 53.8% of urban transient dogs (p=0.012, Table 3). Lost dogs were a more common cause for periurban transient dogs (24.4%) compared to urban transient areas (7.7%) (Table 4), while dogs given away as gifts was a more common cause for urban transient dogs (26.9%) compared to periurban transient areas (4.4%).

We observed a rapid decrease in protection against rabies after the MDVC 2016 campaign until before the following MDVC in 2017 as an effect of births, deaths, and transient dogs. In general, the decline in coverage was more pronounced in periurban areas compared to urban areas. When only births and deaths were considered, in the periurban area, vaccination coverage fell from 68.3% to 34.8% and in the urban area from 60.4% to 38.9% between MDVCs campaigns in 2016 and 2017. We observed that the presence of *transient dogs* negatively affects coverage in both areas, but the impact was more significant in periurban areas. When *transient dogs* were included in the analysis, vaccine coverage decreased to 24.5% in the periurban area and to 34.8% in the urban area (Figure 3). We assumed that the *transient dogs* were not vaccinated and their presence was 3 months on average between the period MDVC 2016 and MDVC 2017.

**Figure 3.**
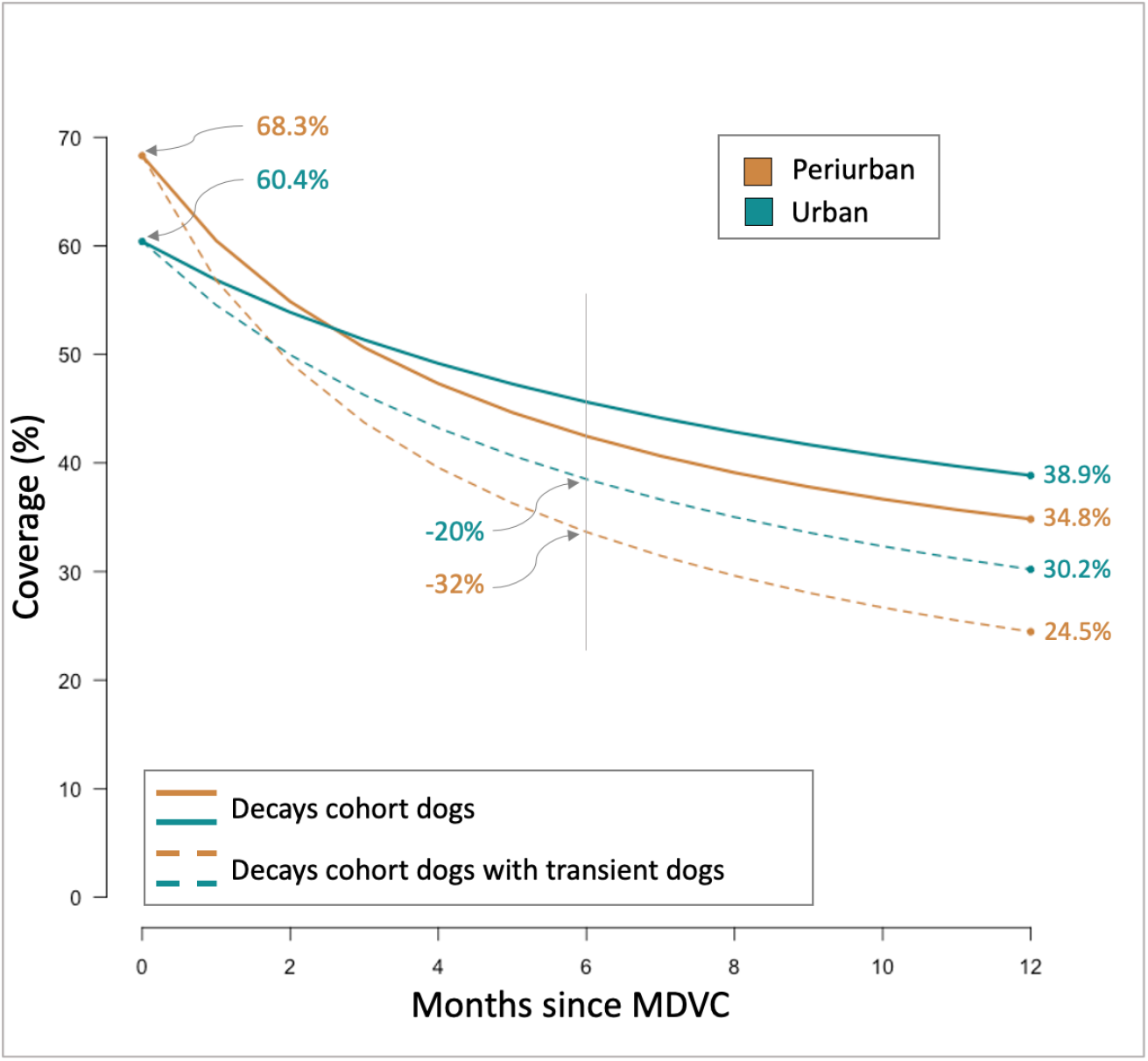
Effect of canine population dynamics on effective vaccine coverage in Arequipa, Peru, estimated with logistic regression models.

During the study period, 248 dogs entered the cohort; 79 into the periurban area and 169 into the urban area (Table 2). New dogs were most commonly obtained as a gift to the owner, especially in periurban areas (Table 2). Breeding of current household dogs was other important contributor to growth of the dog population in both areas. Dog purchases from pet stores or from an acquaintance represented about 11% of new dogs in both areas. Dogs bought from a friend, adopted from a shelter, and other sources all constituted less than 5% of dog acquisitions in either region and these results did not vary by year of study (Table 2). Importantly, in urban areas over 60% of new dogs were sourced from other districts, compared to only 36% in periurban areas. This difference likely reflects the established wide social networks of urban dwellers as opposed to new immigrants found in periurban areas. We also studied dogs that left the cohort. Dog emigration was also associated with the area: while all dogs that emigrated from the periurban area moved only to other districts within Arequipa, 39% of urban dogs moved outside of Arequipa, like Lima (16.7%), or other regions (22.2%) including Cuzco and Moquegua.

**Table 2.** Origin of dogs that entered the cohort.

| Variables | Periurban (n = 79) | Urban (n = 169) | p |
| --- | --- | --- | --- |
| Mode of obtaining |  |  |  |
| Gift | 55.7 | 40.8 | 0.220<sup>a</sup> |
| My dog's offspring, born here | 20.3 | 23.1 |  |
| Bought from store | 10.1 | 12.4 |  |
| Bought from a friend | 2.5 | 5.3 |  |
| Adopted from the street | 11.4 | 12.4 |  |
| Adopted from a shelter | 0.0 | 1.2 |  |
| Other | 0.0 | 4.7 |  |
| Place of obtaining |  |  |  |
| Another house in the neighborhood | 62.0 | 36.1 | 0.002<sup>a</sup> |
| Another District | 13.9 | 31.4 |  |
| Another area of the city | 2.5 | 3.0 |  |
| Another department | 1.3 | 1.8 |  |
| Born here | 20.3 | 27.8 |  |
| p value estimated with (a) Fisher test |  |  |  |

p value estimated with (a) Fisher test

### Dog restriction level across the urban environment

The level of restriction imposed on domestic dogs is an important variable both for understanding dog population dynamics and the transmission of rabies virus and other pathogens. Dogs from the periurban area were found to have greater unsupervised access to the street (p = 0.115) compared to dogs from the urban areas (p <0.001) (Table 3). A greater proportion of dogs in the urban area were taken out for walks by their owners on the street, with a reduction observed from the first year to the second year across both study sites (p <0.001) (Table 3). Similarly, owners in the urban area were more likely to walk their dog on a leash (Table 3).

**Table 3.** Level of dog street access.

| Number of dogs | Periurban |  |  | Urban |  |  |
| --- | --- | --- | --- | --- | --- | --- |
|  | 2016<br>(n=341) | 2017<br>(n=370) | p | 2016<br>(n=956) | 2017<br>(n=1018) | p |
| Free circulation to the street at some time during the day (%)* | 53.1* | 59.8 | 0.115<sup>a</sup> | 20.0* | 37.6 | < 0.001<sup>a</sup> |
| Restriction level (%) |  |  |  |  |  |  |
| Confined / Restricted |  | 38.8 |  |  | 64.0 | <0.001<sup>b</sup> |
| Sometimes it escapes the house) |  | 20.4 |  |  | 14.9 |  |
| Leaves house once or twice a day |  | 16.7 |  |  | 10.3 |  |
| Free circulation to the street |  | 24.1 |  |  | 10.7 |  |
| Owners take the dog for a walk (%) | 48.3 | 27.3 | <0.001<sup>a</sup> | 71.9 | 47.9 | < 0.001<sup>a</sup> |
| Use of leash when walking (%) | 30.8 | 48.0 | 0.012<sup>a</sup> | 49.5 | 76.7 | < 0.001<sup>a</sup> |

(a) Chi square test; (b) Fisher test; \* question was refined in year 2.

**Table 4.**
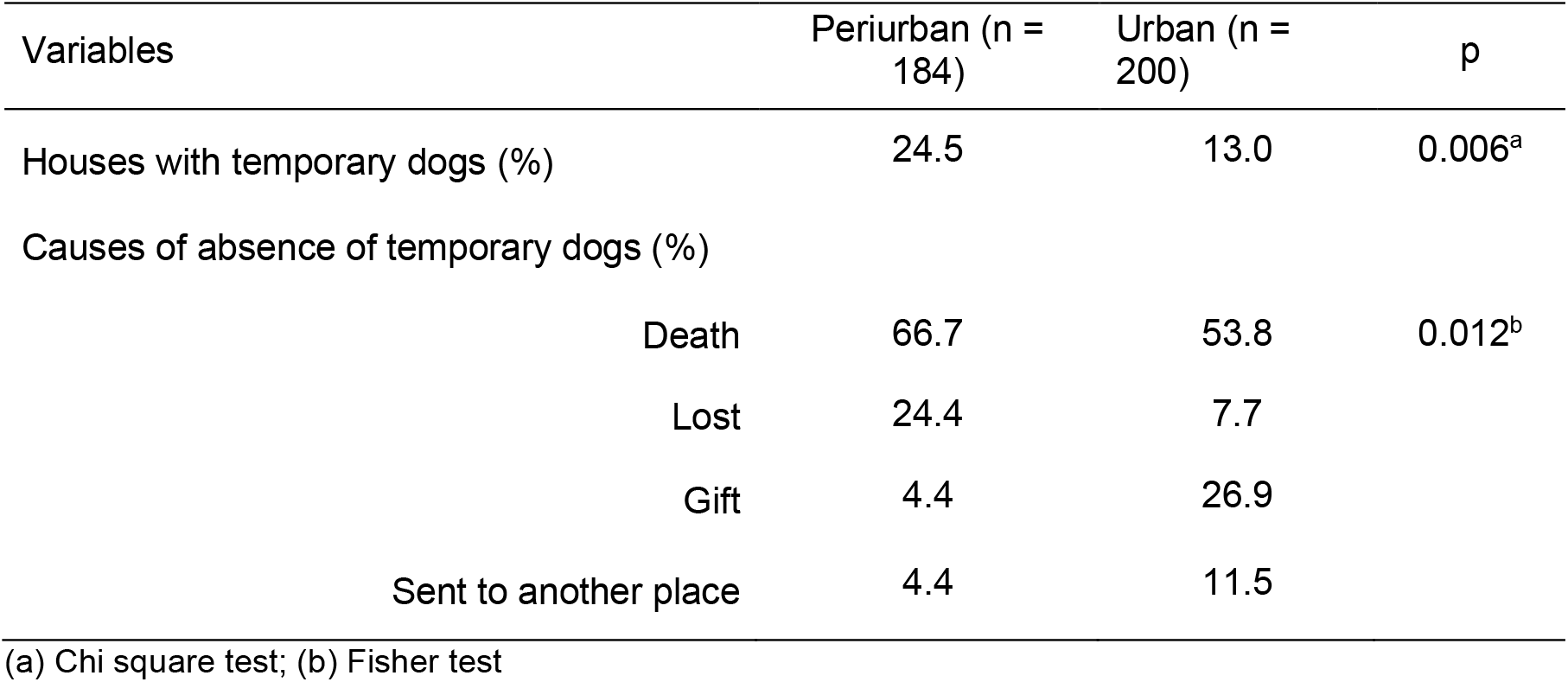
*Transient dogs* in Arequipa, Peru.

| Variables | Periurban (n = 184) | Urban (n = 200) | p |
| --- | --- | --- | --- |
| Houses with temporary dogs (%) | 24.5 | 13.0 | 0.006<sup>a</sup> |
| Causes of absence of temporary dogs (%) |  |  |  |
| Death | 66.7 | 53.8 | 0.012<sup>b</sup> |
| Lost | 24.4 | 7.7 |  |
| Gift | 4.4 | 26.9 |  |
| Sent to another place | 4.4 | 11.5 |  |

(a) Chi square test; (b) Fisher test

### Female dogs with offspring decreased significantly throughout the study

We observed a drastic statistically significant reduction of females with offspring between study years 1 and 2. In the periurban areas, females with offspring went from 32.63% to 9.57% (Chi2 = 15.89; p<0.001) and in urban areas this estimate went from 20.11% to 11.9% (Chi2 = 8.60; p=0.003). Similarly, the yearly average number of puppies per female in the periurban area varied from 1.46 to 0.49 (Poisson distribution; p=0.012) while in the urban area it varied from 0.85 to 0.53 (Poisson distribution; p=0.086). The mean number of puppies per calved female was slightly higher in both years in the periurban area (4.48 and 5.09) compared to both years in the urban area (4.23 and 4.43).

### Dog population dynamics and rabies immunity

We built a basic rabies virus transmission model to evaluate the effect of the extremely high population turnover rates on the numbers of rabid dogs in Arequipa. An SEIV system was simulated over a period of 900 days. The model was parameterized with different death rates for different areas (Table 5); birth rates were subsequently derived to keep population levels in equilibrium. The resulting simulations (Figure 4) show how higher population turnover in periurban communities can theoretically drive higher dog rabies incidence.

**Table 5.** Dogs exiting the cohort in Arequipa, Peru.

| Variables | Periurban (n = 133) | Urban (n = 268) | p |
| --- | --- | --- | --- |
| Female (%) | 30.1 | 33.6 | 0.553<sup>a</sup> |
| Age of dogs in months - Median (IQR) | 24 (18 - 48) | 36 (24 - 72) | < 0.001<sup>b</sup> |
| Age of dogs in months - Mean (SD) | 43.68 (39.09) | 56.4 (45.6) | 0.004<sup>c</sup> |
| Purebred (%) | 6.8 | 22.4 | <0.001<sup>a</sup> |
| Vaccinated (%) | 7.1 | 23.6 | 0.058<sup>a</sup> |
| Free access to the street (%) | 54.1 | 19.8 | < 0.001<sup>a</sup> |
| Causes of dogs that left the cohort (%) |  |  |  |
| Death | 86 | 75 | 0.276<sup>d</sup> |
| Send to another place | 10.5 | 17 |  |
| Lost | 3.5 | 8 |  |
| Emigration of dogs (%) |  |  |  |
| To other districts within Arequipa | 100 | 61.1 | 0.36<sup>d</sup> |
| To Lima | 0 | 16.7 |  |
| To other regions | 0 | 22.2 |  |
| Causes of deaths (%) |  |  |  |
| Poison | 40.8 | 16.7 | < 0.001<sup>d</sup> |
| Disease | 4.1 | 34.5 |  |
| Run over | 24.5 | 20.2 |  |
| Natural/Old age | 8.2 | 25 |  |
| Other | 22.4 | 3.6 |  |

(a) Chi square test; (b) Test of Mann-Whitney-Wilcoxon, (c) Student t-test, (d) Fisher test

**Figure 4:**
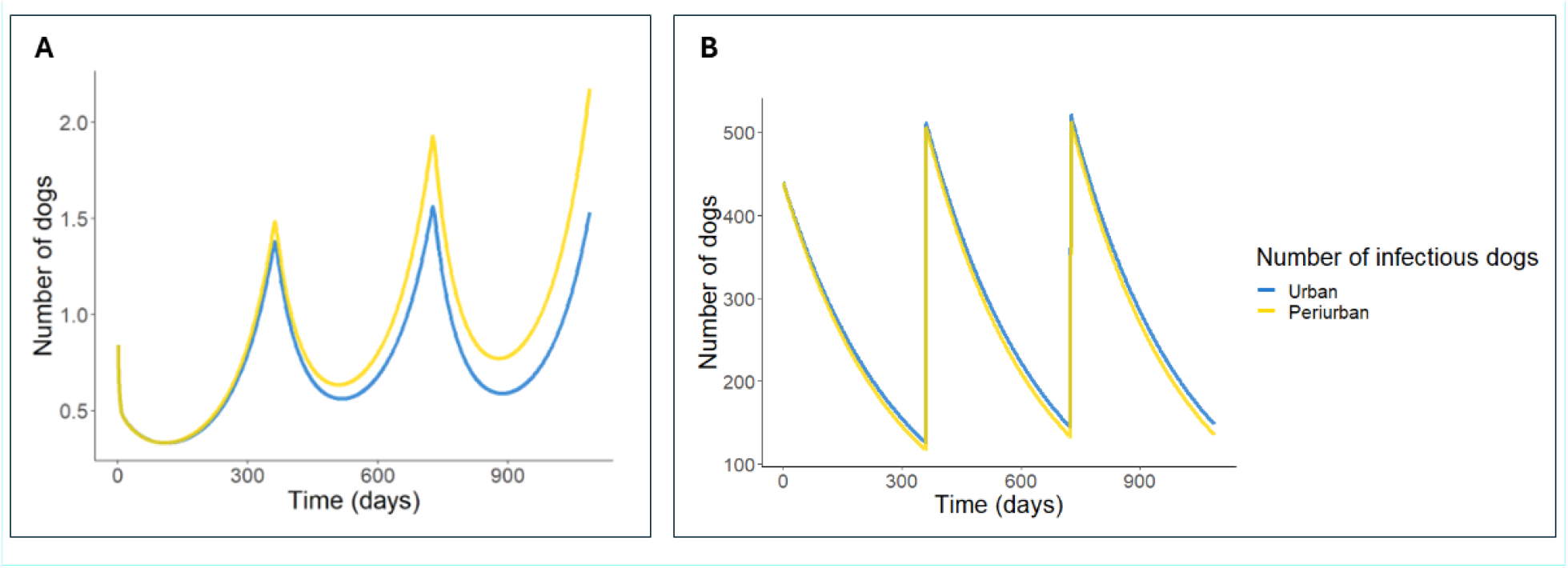
Three years simulation of canine rabies in Arequipa, Peru, assuming equivalent vaccination rates. Panel A indicates simulated number of infections while Panel B indicates population vaccination levels.

## DISCUSSION

We conducted a 12-month survey over 04 urban and 06 periurban communities to study the population dynamics of owned dogs in a dog rabies-endemic area in southern Peru. We found a highly dynamic population, with rapid turnover and growth, and with significant variability in demographic parameters across geographic areas. Authorities are struggling to eliminate dog rabies from this population, primarily due to suboptimal annual vaccination coverage. Furthermore, as our study suggests, the ever-changing dynamics of these dog populations make it difficult to maintain current parameter estimates to inform control efforts, making it harder to achieve rabies elimination. Using an outdated human to dog ratio, in 2015, health authorities reported a rabies vaccination coverage of 100% in the dog population of Arequipa^26^. We found that the human to dog ratios were significantly lower than the ratio established by the MOH in the MDVC (∼4:1 vs 6:1)^17,25^, which can lead to severe underestimations of the actual dog population creating the illusion of achieved herd immunity when coverage levels are in fact suboptimal^16^.

The human to dog ratios varied significantly by geographical area, with a lower ratio in the periurban localities compared to urban localities. In the rapidly expanding periurban areas, there is more uncontrolled breeding as resident free-roaming owned dogs encounter and absorb migrant populations^26,30^. Combined with our findings on dog movement in periurban areas^5^, these human to dog ratios highlight periurban areas as key geographic targets for future interventions. Our estimated =ratios align with those reported in other studies in Arequipa for both periurban areas and urban areas^31–34^, suggesting these findings may be generalizable to other parts of the city. Historically, rabies control reports in Latin America have used a standard 10:1 human to dog ratio for countries ranging from Argentina and Venezuela, to Haiti and Cuba^35^. However, ratios in other Latin American countries are often much lower and more similar to or slightly higher than those estimated for our urban areas, ranging from 4.0:1 to 4.8:1 in Mexico^36^, Bolivia, Brazil, and Chile^37–40^. Globally, these ratios vary widely: in Africa, mean ratios are 21.2:1 for urban areas and 7.4:1 for rural areas, while in Asia, they are 7.5:1 for urban and 14.3:1 for rural areas^41^.

The movement of people with animals between Arequipa and the Puno region occurs due to the abundance of temporary jobs in agriculture and mining in Arequipa^42^. Genetic sequencing has shown that the rabies virus circulating in Arequipa was originally introduced from Puno, signaling the reintroduction of the virus to a nonendemic region^43^. In this study, we found dogs being brought to Arequipa from several other regions including Cuzco, Abancay, Lima and Puno, which could potentially produce new reintroductions of rabies virus into the city of Arequipa. We also found the exportation of dogs from this rabies-affected area to regions like Cuzco, Moquegua, and the capital Lima. the latter being a large city of 11 million people that is currently free of rabies. However, Lima’s rabies-free status might be threatened by current evidence suggesting insufficient population immunity and potential gaps in surveillance, possibly resulting from inadequate vaccination coverage in some areas^44^. Almost all dogs that were relocated to Cuzco, Moquegua, and Lima were from the urban areas which may be related to the socioeconomic means of the residents, with those who can afford to live in the more expensive urban localities able to relocate farther. Therefore, the constant relocation of dogs without evaluation of vaccination status, has serious implications for local transmission of rabies virus and spread of rabies into new rabies-free areas, potentially putting larger populations at risk^45–47^.

The dog population is growing rapidly in Arequipa, and our data indicate that local reproduction is more important than immigration in explaining this growth. Principally responsible for this increase is the raising of reproductively active females on the street; up to 55.2% of female dogs in the periurban area, and 9.4% of female dogs in the urban area are reproductively active. Multiple authors have reported a clear preference for owning male dogs over female dogs^38,48,49^ as they are considered better guardians and hunters^30,50^. Our study corroborates this preference, as six to seven out of every ten registered dogs were male, with a higher proportion in the periurban area^38,51–54^. Females are seen as a problem for attracting males during estrus^55^ and as a result, the majority of unwanted female offspring are given away, sold, euthanized, or abandoned on the street^3^. Females that are lost or abandoned on the street can contribute several litters per year impacting population control programs^56^. Moreover, free-roaming dogs are generally poorly fed and do not receive veterinary attention. So they move around frequently in search of food from garbage dumps^19^, increasing the interactions between dogs and facilitating transmission of the rabies virus^30^. This combination of uncontrolled breeding and increased contact between individuals has serious impacts on rabies virus transmission and vaccine coverage.

Like births, dog deaths also influence effective vaccination coverage. We found a striking difference in dog mortality causes between urban and periurban areas; the periurban area registered four times more deaths from poisoning compared to the urban area. Dog culling, which has been implemented in some areas as a rabies control measure, may increase the dog population turnover as people rapidly replace the culled dog (possibly immunized against rabies) with a new susceptible dog (from the same district or another area) causing a reduction in herd immunity^57^. Deaths from rabies and other diseases also accelerates the turnover rate of dogs.

For example, in some areas, canine parvovirus (CPV-2) kills up to 90% of infected young dogs^58^ and *Echinococcus granulosus* infections pose an increasing threat to the longevity of dogs^22^. Our study and others’ suggest that one of the main barriers to any successful dog rabies vaccination program in the developing world is the high proportion of very young, unvaccinated dogs^4^. This dynamic of rapid turnover in the dog population generates a positive feedback loop, as the high death rate causes a reduction in immunity which leads to further death as a result of rabies disease, increasing the need for more frequent vaccination campaigns, especially in periurban areas^51^.

A unique aspect of our study was the inclusion of *transient dogs* as a variable of interest. *Transient dogs* are individuals that entered and exited the cohort in the time between survey years, which otherwise would have been unaccounted for. We found that 27% of households in the periurban areas and 17% of households in the urban areas housed a *transient dog*. Furthermore, the majority of these dogs were lost as a result of death (66.7% in periurban localities; 53.8% in urban localities). *Transient dogs* represent a population with extremely high turnover (less than one year), indicating an important drop in vaccination coverage that has gone undetected. Using a basic SEIV mathematical model, we demonstrate how higher turnover of the dog population can lead to increased rabies incidence. This occurs because vaccinated dogs are replaced with susceptible puppies increasing the pool of susceptible individuals. This correlates with studies worldwide suggesting that high dog population turnover can drive rabies persistence^3,4,37,38,49,50,54,59–65^. Large urban areas of Peru, with likely similar dog population dynamics, are affected by other dog-associated zoonosis including Chagas disease^21^, echinococcosis^22^, and bartonellosis^66^. Highly dynamic dog populations could similarly affect public health programs to control these zoonotic diseases.

This study had several limitations. Data were collected by household surveys and in some instances the family members available were children or elderly individuals who could not be adequately questioned due to their age or lack of understanding. As with any survey data, there is also the possibility of memory and information bias from participants, and it is possible that the data received from households was not entirely accurate. For example, it was often quite difficult to get precise estimates on the dogs’ ages. A major strength of this work is the inclusion of *transient dogs* as a variable of interest. We discovered a large population of these dogs, with major implications for rabies vaccination coverage of the dog population as a whole. We strongly suggest the inclusion of *transient dogs* to any longitudinal study of dog borne zoonoses as they may represent an area of risk previously unaccounted for.

Since domestic dogs are the main reservoir of rabies virus, controlling the disease lies in recognizing the implications of the demographic characteristics of dog populations and informing control measures accordingly^59^. We found rapid canine population growth (an 8.5% increase in periurban areas and 7.5% increase in urban areas) and a very high population turnover (48% in periurban areas and 34% in urban areas). These two parameters cause a rapid decrease in effective vaccination coverage between MDVCs, especially in periuban areas. Population turnover is an important threat to achieving the minimum vaccine coverage established by the WHO (70%)^67^ and the Pan American Health Organization (80%)^68^. Prevention programs for rabies and other dog-associated zoonoses must consider these dynamics in their planning. Addressing population turnover will require coordinated planning, dog owner education, and comprehensive veterinary medical care to treat dog diseases to prevent excessive dog deaths. The “hotspotting” strategy in human health focuses preventive health resources on people with multiple comorbidities to improve outcomes and reduce health spending^69,70^. Our findings point to periurban areas as a “hotspot” for targeted health interventions. These should include vaccination, veterinary assistance, and education for residents about responsible dog ownership behavior and the transmission of dog-borne diseases. Effective control of rabies and other zoonotic diseases requires a holistic strategy that addresses the health of domestic dogs and their impact on human health. By understanding and managing the dynamics of dog populations, we can better protect both human and animal health.

